# ALI MEA60: A porous thin-film microelectrode array for air–liquid interface electrophysiology of 3D excitable tissues

**DOI:** 10.64898/2026.11.16.766976

**Authors:** Fabien Friscourt, Luc Stoppini, Shaginth Sivakumar, Marc O. Heuschkel, Laetitia Nikles, Théo Ribierre, Thi-Thom Nguyen, Michel Pucéat, Marisa Jaconi, Léonardo Scapozza, Adam Armada-Moreira, Laura Monni, Alessio Di Clemente, Michele Giugliano, Julio Alvarez-Collazo, Jannis E. Meents, Sven Schoenecker, Adrien Roux

## Abstract

**Objective:** Conventional microelectrode arrays (MEAs) are suboptimal for long-term electrophysiology of three-dimensional (3D) excitable tissues, as limited oxygenation, nutrient diffusion, and tissue-electrode coupling compromise recording stability. Air-liquid interface (ALI) culture mitigates these constraints, but existing ALI MEA platforms are limited in channel count, flexibility, and compatibility with commercial multichannel acquisition systems. We developed and comprehensively characterized a next-generation ALI MEA for standardized, long-term recordings across multiple 3D excitable tissue models.

**Approach:** The novel ALI MEA60 is composed of a porous polyimide membrane that integrates an array of 60 thin film microelectrodes (59 recording, one reference) and a basal perfusion chamber with Luer-lock connectors for controlled medium exchange and pharmacological wash-in/wash-out. It is compatible with the MEA2100-Mini data acquisition system (Multi Channel Systems). We characterized electrode impedance and noise across 41 ALI MEA60 devices, assessed 60-day stability with human iPSC-derived neural spheroids (N = 7 MEAs, 121 recordings) using a standardized longitudinal framework and linear mixed-effects models, and evaluated its versatility through pharmacological assays and recordings from human neural spheroids, brain organoids, ex vivo brain slices, and cardiac organoids.

**Main results:** The platform showed reproducible electrical performance (median impedance 56 kΩ at 1 kHz) and low noise (≈5 µV Root Mean Square (RMS) in saline, 10–20 µV in medium), resolving spikes and local field potentials simultaneously. Recording remained stable for at least 60 days, with a stable noise floor and ∼85% persistence of initial functional channels. Neural spheroids reproduced the standard qualitative signatures to gabazine, 4-aminopyridine (4-AP), and 6-Cyano-7-nitroquinoxaline-2,3-dione/D-2-amino-5-phosphonovalerate (CNQX/D-AP5), while cardiac organoids showed the expected β-adrenergic response to isoproterenol. The ALI MEA60 can similarly record signals from human brain slices and organoids without hardware modification.

**Significance:** The ALI MEA60 combines ALI culture, chronic recording stability, standardized pharmacological testing and cross-tissue compatibility in a single ready-to-use platform. It also establishes a reproducible framework for benchmarking long-term electrophysiological performance, thereby supporting disease modelling, drug discovery, organ-on-chip technologies and future chronic 3D electrophysiology systems.

## 1 INTRODUCTION

Three-dimensional (3D) cell culture models including spheroids, organoids, and assembloids are widely used for disease modelling, drug screening and developmental biology, as they reproduce native tissue architecture, cell–cell interactions and aspects of *in vivo* function more faithfully than two-dimensional (2D) cultures ^1–3^. In neurodevelopment and neuronal disease research, organoids and assembloids enable investigation of complex neural connectivity and physiological activity, while cardiac organoids are also increasingly used for electrophysiological and pharmacological testing ^3–5^. Compared with monolayer cultures, organoid systems better preserve parental tissue features and support longer-term maintenance of biological characteristics, thus increasing the demand for functional assays that remain stable over extended culture periods ^1,6^. Beyond stem-cell-derived models, *ex vivo* preparations such as acute brain slices and *ex vivo* explants remain reference systems for studying native neural circuits with their cytoarchitecture and local connectivity preserved. Between acute slices and stem-cell-derived 3D systems lie organotypic slice cultures, i.e. thin explants maintained for weeks at the air–liquid interface on a porous membrane^7^, as a long-standing *ex vivo* model that bridges the two and is naturally supported by the same interface. Like acute slices, these preparations have a shorter viability window than 3D cultures but share the need for recording interfaces that access thick, non-adherent tissue rather than a planar monolayer.

Microelectrode arrays (MEAs) are an established label-free, non-invasive technology for extracellular recording of excitable tissues *in vitro* ^6,8,9^. Originally, most conventional MEAs are built as planar electrode grids on rigid glass substrates, optimised for 2D cultures in which every cell lies directly on the recording plane. When such a rigid, closed-bottom substrate holds a 3D tissue submerged in medium, only the basal face contacts the electrodes, and the tissue depends on diffusion through a bulk-medium column for oxygen and nutrients, a configuration that limits both spatial coverage and long-term viability, and promotes tissue flattening ^5,8–10^. This geometric mismatch becomes problematic for spheroids and organoids larger than ∼500 µm^11,12^, where limited oxygen and nutrient diffusion, restricted solution exchange at the electrode-contacting face, and weak or unstable tissue–electrode coupling can compromise viability and signal stability during multi-week experiments ^9,13^.

Recent 3D-interfacing MEA platforms such as high-density complementary metal oxide semi-conductor (CMOS) or 3D geometry (i.e., mesh, shell, stretchable, pillar, pyramid, and wrapping) designs address this limitation by increasing electrode density, surrounding the tissue, or penetrating toward inner cell layers ^9,14^. These systems can improve signal quality and spatial coverage, but often introduce trade-offs in handling complexity, predefined geometry, as well as specialised holders, custom fabrication, or hardware integration requirements ^12,15,16^. Several reports also note reliance on continuous perfusion, frequent medium renewal, or other *ad hoc* measures limiting central necrosis and maintaining tissue health during long-term recording ^9,12,17^.

Air-liquid interface (ALI) culture offers a complementary strategy in which 3D tissues are supported on a porous membrane, oxygenated directly through the apical surface, and fed basally through the membrane ^4,18^. Porous-membrane MEAs were originally developed to maintain this interface around 3D neural tissues and support long-lasting electrophysiological monitoring without active perfusion of the tissue bulk, with reported experiments extending over several weeks ^17,18^. Beyond 3D cultures, this interface has long been used to preserve nervous-tissue slice viability for extracellular and patch-clamp recordings^7^, connecting the present platform to established slice electrophysiology. Building on this lineage, current efforts position porous-membrane MEAs toward higher channel counts, flexible tissue-placement configurations and improved compatibility with modern acquisition workflows ^16,18^.

In this paper, we describe the ALI MEA60, a new-generation porous polyimide substrate-based thin-film microelectrode array device that extends the ALI MEA lineage along three practical axes within a single, standardised device: 1) scaling to 59 recording electrodes and one reference electrode on an 8 × 8 grid (200 µm pitch, 30 µm platinum (Pt)-black-coated sites); 2) integration of a basal perfusion chamber accessible through standard Luer-lock connectors for reproducible medium exchange and pharmacological wash-in/wash-out; and 3) direct compatibility with the MEA2100-Mini data acquisition system (Multi Channel Systems MCS GmbH). Beyond the device itself, we introduce a standardised longitudinal framework that extracts 32 recording-level metrics, 12 of them reported here as representative, spanning signal quality, channel yield and network activity across MEAs over 60 days. This provides a transparent, quantitative basis for reporting the platform’s stability, benchmarking it against alternatives, and guiding future iterations.

Ultimately, this work characterizes the platform as a tool - its specifications, the signals it captures reliably, and the assays it enables - for users considering adoption in their own 3D-tissue electrophysiology workflows. We illustrate this on four different biological models: human iPSC-derived neural spheroids and organoids, human cardiac organoids, and *ex vivo* human brain slices, which together cover the representative use cases of long-term extracellular electrophysiology.

## 2 METHODS

### 2.1 Device design and fabrication

The ALI MEA60 is a 60-channel porous polyimide substrate-based thin-film MEA designed for ALI culture of 3D excitable tissues and compatible with the MEA2100-Mini acquisition system (Multi Channel Systems MCS GmbH). The device integrates 59 recording/stimulation electrodes and one reference electrode in an 8 × 8 grid (corner positions omitted) on an 8 µm-thick polyimide membrane. Recording sites are 30 µm in diameter and arranged on a 200 µm grid. Electrodes are coated with Pt-black to reduce impedance and increase effective charge-transfer capacity. The polyimide membrane is perforated with through-holes (7.5 µm diameter, 20 µm pitch), yielding an open porosity of approximately 10% across the active area. The porous geometry allows basal medium to reach the underside of the tissue while the apical surface remains exposed to humidified incubator air, establishing ALI conditions across the active area.

The membrane is integrated into a custom perfusion chamber housing mounted on a printed circuit board (PCB) that provides electrical interfacing with the data acquisition system. A basal chamber of 350 µL volume located under the porous membrane is accessed through two Luer-lock connectors, enabling either static culture or controlled medium exchange and pharmacological wash-in/wash-out. A 0.45 µm hydrophilic PTFE membrane is incorporated within the fluidic path to homogenize basal flow and limit bubble formation. The device is mechanically sealed for leak-free operation under standard incubator conditions. Photolithographic patterning, thin-film metallization, Pt-black electrodeposition, perforation, and bonding / sealing procedures are described in detail elsewhere^19^.

### 2.2 Electrical characterization

Electrode impedance and phase were measured at 1 kHz in physiological saline (0.9% NaCl) at room temperature using the MEA-IT instrument (Multi Channel Systems), after Pt-black electrodeposition and prior to tissue placement (N = 41 MEAs, 59 electrodes per chip). The impedance of the electrodes was measured relative to an external Ag/AgCl reference electrode. Baseline noise was quantified as the RMS amplitude of raw signals over 10s windows under cell-free saline and culture-medium conditions, and during representative chronic culture recordings.

### 2.3 Air-liquid interface culture on the device

Three-dimensional tissues were placed directly on the porous electrode area. The basal chamber was filled with the appropriate culture medium via the Luer-lock ports, and the apical surface was left exposed to humidified incubator air (37 °C, 5 % CO_2_). No coating or adhesion promoter was used as it is not necessary to achieve good tissue/device coupling. Medium exchange through syringe use was performed through the basal Luer-lock ports using two sequential exchanges of the basal volume to improve wash-in/wash-out efficiency in the low-volume chamber.

### 2.4 Tissue preparation

#### 2.4.1 Human neural spheroids

Human neural spheroids were generated from human iPSC-derived neural stem cells (NSChiPS, Thermo Fisher Scientific) following our previously published protocol^20^ with minor adaptations. NSChiPS were expanded on GelTrex-coated flasks in KnockOut™ DMEM⁄F-12 Basal Medium (Thermo Fisher Scientific) supplemented with Gibco StemPro NSC SFM (Thermo Fisher Scientific), Epidermal Growth Factor (EGF) and Fibroblast Growth Factor 2 (FGF-2) to ∼80% confluency, then dissociated with Accutase and seeded at 2.5 × 10⁵ cells/well in 6-well plates under orbital shaking (80 rpm, 37 °C, 5% CO_2_). Neural differentiation followed a two-step medium transition. DIFF1 (per 50 mL: 44.45 mL DMEM/F-12 with GlutaMAX, 3.6 mL 25% BSA, 1 mL StemPro hESC Supplement, and 100 µL BDNF, 100 µL GDNF, 250 µL dibutyryl-cAMP and 500 µL 2-phospho-ascorbic acid stock solutions) was used during week 1, and DIFF2 (a 1:1 mix of DIFF1 and NDM) during weeks 2–4, as previously published²⁰. From week 5 onward, spheroids were maintained in NDM (Neurobasal™ Plus supplemented with B27™ Plus and GlutaMAX™) with weekly medium changes^20^. Spheroids reached an average diameter of 500–600 µm and developed spontaneous network activity from weeks 6-8 onward. Prior to ALI MEA experiments, between 4-6 months old spheroids were pre-conditioned for 24-48 h under standard ALI conditions on pre-cut patches of hydrophilic PTFE membranes placed on culture inserts. Only spheroids displaying preserved structural integrity and homogeneous optical density were used for electrophysiological recordings.

#### 2.4.2 Human cardiac organoids

Human cardiac organoids were generated according to a protocol modified from Hofbauer et al. ^21^. Briefly, 2,500 H9 TnT-GFP human embryonic stem cells (WiCell) were aggregated in StemFit4 medium (Amsbio). After 30 h, aggregates were induced with 1 mM CHIR99021, 10 ng/mL BMP4 and 75 ng/mL activin A in cardiac differentiation medium (CDM) for 36 h. CDM consisted of F-12 (Ham) with GlutaMAX and IMDM with GlutaMAX (1:1) supplemented with BSA fraction V (5 mg/mL), thioglycerol (450 µM), insulin (7 µg/mL), transferrin (15 µg/mL) and chemically defined lipid concentrate (1×). Cells were then treated with 10 ng/mL BMP4, 8 ng/mL FGF2, 0.5 mM retinoic acid and 5 mM XAV939 (MedChemExpress) in CDM plus insulin for 4 days, followed by BMP4 and FGF2 only in CDM plus insulin for 2 days. Beating organoids were maintained in CDM with insulin for two further weeks. Mature, spontaneously beating organoids (15 DIV) were transferred onto the ALI MEA in CDM.

#### 2.4.3 Human brain organoids

Commercially available healthy female patient-derived hiPSC purchased from Phenocell were used for the generation of dorsal cortical brain organoids following a previously published protocol^22^. Briefly, 2 × 10⁶ hiPSC were aggregated on day 0 and sequentially exposed to cocktails of patterning molecules (days 0–6: DM, SB431542, XAV939; days 7–24: EGF, FGF; days 25–42: BDNF, NT3) over 43 days, then cultured under agitation in a maturation medium consisting of Neurobasal and SM1-A. Here, around 4-months-old organoid were used.

#### 2.4.4 Ex vivo human brain slices

Human cortical tissue obtained from patients undergoing resective surgery for pharmacoresistant epilepsy was processed as follows. Immediately after surgical resection, tissue was transported from the operating room to the laboratory within 30 min in an ice-cold, carbogenated (95% O_2_/5% CO_2_), sterile-filtered NMDG-based artificial cerebrospinal fluid (aCSF) containing (in mM): NMDG 92, HEPES 20, Na-pyruvate 3, taurine 0.01, thiourea 2, glucose 25, myo-inositol 3, NaHCO_3_ 30, CaCl2 0.5, MgSO_4_ 10, KCl 2.5, NaH_2_PO_4_ 1.25, HCl 92, L-ascorbic acid 5, and N-acetyl-L-cysteine 12. After removal of the pia mater, tissue blocks were sectioned into 350-µm-thick coronal slices using a vibratome in ice-cold carbogenated NMDG-based aCSF. Slices containing the full cortical thickness (layers I–VI) together with the underlying white matter were incubated for 15 min at 34 °C in the same solution and subsequently transferred to a Na-based recovery aCSF containing (in mM): NaCl 92, HEPES 20, Na-pyruvate 3, taurine 0.01, thiourea 2, glucose 25, myo-inositol 3, NaHCO_3_ 30, CaCl_2_ 2, MgSO_4_ 2, KCl 2.5, NaH_2_PO_4_ 1.25, L-ascorbic acid 5, and N-acetyl-L-cysteine 12, for 30 min at room temperature. Following recovery, slices were mounted onto the ALI-MEA containing pre-warmed culture medium and maintained at 37 °C in a humidified incubator (5% CO_2_) until electrophysiological recordings. The culture medium consisted of Minimum Essential Medium (MEM; Gibco 21090022) supplemented with (in mM): L-ascorbic acid 1, CaCl_2_ 1, MgSO_4_ 2, HEPES 30, glucose 13, Tris base 1, and glutamine 2.5, together with 20% heat-inactivated horse serum, 0.2% penicillin/streptomycin, and bovine pancreas insulin (1 mg/l). All solutions were adjusted to pH 7.3–7.4, an osmolarity of 300 ± 5 mOsm/kg, sterile-filtered before use, and all tissue handling procedures were performed under near-sterile conditions.

### 2.5 Acquisition and signal processing

Recordings were performed on the MEA2100-Mini system at a sampling rate of 25 kHz/channel (10 kHz for cardiac organoid field-potential recordings to reduce file size) and exported in HDF5 format. Channel labels were mapped to electrode coordinates using a ProbeInterface-compatible geometry file^23^.

*Per-channel quality control (QC)* was based on the variance, RMS amplitude, and median absolute deviation (MAD; σ ≈ MAD / 0.6745) of the raw signal over a fixed 30– 60 s window. Channels were flagged as bad and excluded from downstream analyses if their variance fell below 1 × 10^-6^ µV^2^ or if their RMS or MAD exceeded the across-channel median plus 5 × MAD.

#### Multi-unit activity (MUA)

Signals were band-pass filtered (300–3000 Hz, zero-phase 4^th^-order Butterworth) and spikes were detected on each good channel using a robust MAD-based threshold (k = 5.5), with a 1 ms refractory period and ±0.5 ms peak realignment.

#### Spike sorting

For figures showing per-electrode units (Figure 2(d)), waveforms were extracted in a −1 / +1.5 ms window around the detected peak and decomposed by Principal Component Analysis (PCA, 3 components). Putative units were identified by Gaussian Mixture Model clustering, with the number of components selected by minimum Bayesian Information Criterion (BIC) and capped at 4 units per polarity. Negative- and positive-polarity components were sorted independently.

#### Population rate and network bursts

Population firing rate was obtained by pooling spikes from good channels into 10 ms bins and smoothing with a Gaussian kernel (σ = 30 ms). *Network bursts were identified from single-channel burst* co-occurrence: a network burst was defined as any period during which at least 25% of active electrodes were simultaneously undergoing a single-channel burst (ISI-based, ≥5 spikes, max ISI 100 ms), with merging of segments separated by <150 ms and exclusion of events <50 ms. For each network burst, onset/offset, peak rate, recruited electrode fraction, and per-electrode first-spike latency were extracted. Functional connectivity was quantified using the Spike Time Tiling Coefficient^24^ with a ±20 ms coincidence window.

#### Local field potential (LFP) analysis

Signals were median-centred, band-pass filtered (1–200 Hz, 4^th^-order Butterworth, zero-phase), and down-sampled to 1 kHz using polyphase resampling. Power spectral density was estimated by Welch’s method and integrated over canonical bands (δ 1–4, θ 4–8, α 8–13, β 13–30, γ 30–80, high-γ 80–150 Hz).

*Cardiac field potentials* were processed with an automated beat-detection pipeline (band-pass 0.5-100 Hz for display, 0.5-250 Hz for analysis) yielding beat rate, inter-beat interval, and beat-to-beat variability.

All analyses were performed in Python (NumPy, SciPy, statsmodels, matplotlib). The full analysis pipeline and the recording-level master dataset are available from the corresponding author upon reasonable request.

### 2.6 Pharmacological protocol

#### Neural spheroid protocol

Each neural spheroid received a within-sample sequential pharmacology paradigm: Baseline (NDM, 10 min stabilization + 10 min recording) → gabazine (5 µM, two-step exchange, 20 min recording) → washout (NDM, two-step exchange, 20 min recording) → 4-AP (100 µM, 20 min recording) → washout (NDM, 20 min recording) → CNQX (20 µM) + D-AP5 (50 µM) (20 min recording). DMSO concentration was vehicle-matched across conditions and maintained at ≤ 0.1 % (v/v).

#### Cardiac organoid protocol

Each organoid received: Baseline (CDM, 10 min stabilization + 10 min recording) → isoproterenol (100 µM, 10 min wash-in, 20 min recording) → washout (CDM, 10 min recording). DMSO concentration was matched at ≤ 0.1 % (v/v).

### 2.7 Long-term recordings and statistics

Neural spheroids were recorded chronically on N = 7 chips over 60 days, yielding 121 recordings across ∼50 timepoints. For each recording, 32 metrics were extracted at the recording level (one observation per chip per recording day), grouped as hardware quality (noise RMS, MAD, spike amplitude, Signal-to-noise ratio (SNR), channel counts), spatial coverage (active electrodes, persistent good channels, Jaccard / renewal indices vs. Day 0), and network activity (spike, burst, STTC, and LFP metrics).

#### Outlier exclusion

Recordings were flagged per chip using a two-stage criterion combining a robust intra-chip threshold (median + 3 × MAD from Day 7 onward, MAD scaled by 1.4826) and absolute cutoffs (30 µV moderate, 60 µV extreme for noise RMS; 25 / 50 µV for noise MAD). Recordings flagged by either criterion (2 / 121 in the present dataset; both flagged by the intra-chip MAD criterion) were excluded from inferential statistics but retained in the master dataset.

#### Longitudinal statistics

For each metric, a linear mixed-effects (LME) model was fitted at the recording level with the formula metric ∼ day + (1 | chip) — that is, a fixed linear effect of day (the population-level trend over time) and a random intercept per chip (allowing each chip its own baseline while sharing a common slope) — using maximum likelihood (statsmodels mixedlm, BFGS primary optimizer, Nelder–Mead fallback). This structure separates the average temporal trend from stable chip-to-chip differences, so that the day effect is not confounded by baseline variability between chips. Standardized effect sizes were computed as β × SD (day) / SD (metric) to allow cross-metric comparison. LME p-values were not adjusted across metrics. Post-hoc pairwise comparisons versus baseline were corrected using the Holm–Bonferroni method within each metric.

#### Pharmacology statistics

Electrode-level LME models with chip identity as a random intercept and recording as a variance component were used, with Baseline NDM as the reference condition and Holm correction within each model. Bootstrap mean differences (drug − Baseline, 5,000 iterations, percentile method) at the recording-median level are reported alongside the LME results as effect-size summaries. Direction of all LME-significant effects was independently confirmed by Friedman tests at the recording level. Significance thresholds: *p < 0.05, **p < 0.01, ***p < 0.001, ****p < 0.0001.

## 3 RESULTS

### 3.1 Device and electrical characterization of the ALI MEA platform

The ALI MEA60 was designed to address a persistent constraint of conventional MEAs built on rigid, closed-bottom substrates: their limited ability to keep 3D neural tissue viable during chronic recording. The platform integrates three functional elements within a standard 5 × 5 cm chip carrier compatible with the MEA2100-Mini data acquisition system (Multi Channel Systems MCS GmbH, Harvard Bioscience): a porous polyimide substrate-based thin-film microelectrode array, a basal perfusion chamber accessible through two Luer-lock ports, and a poly(methyl-methacrylate) (PMMA) cover lid that maintains the apical tissue surface exposed to incubator humidified air (Figure 1(a)-(c)).

**Figure 1:**
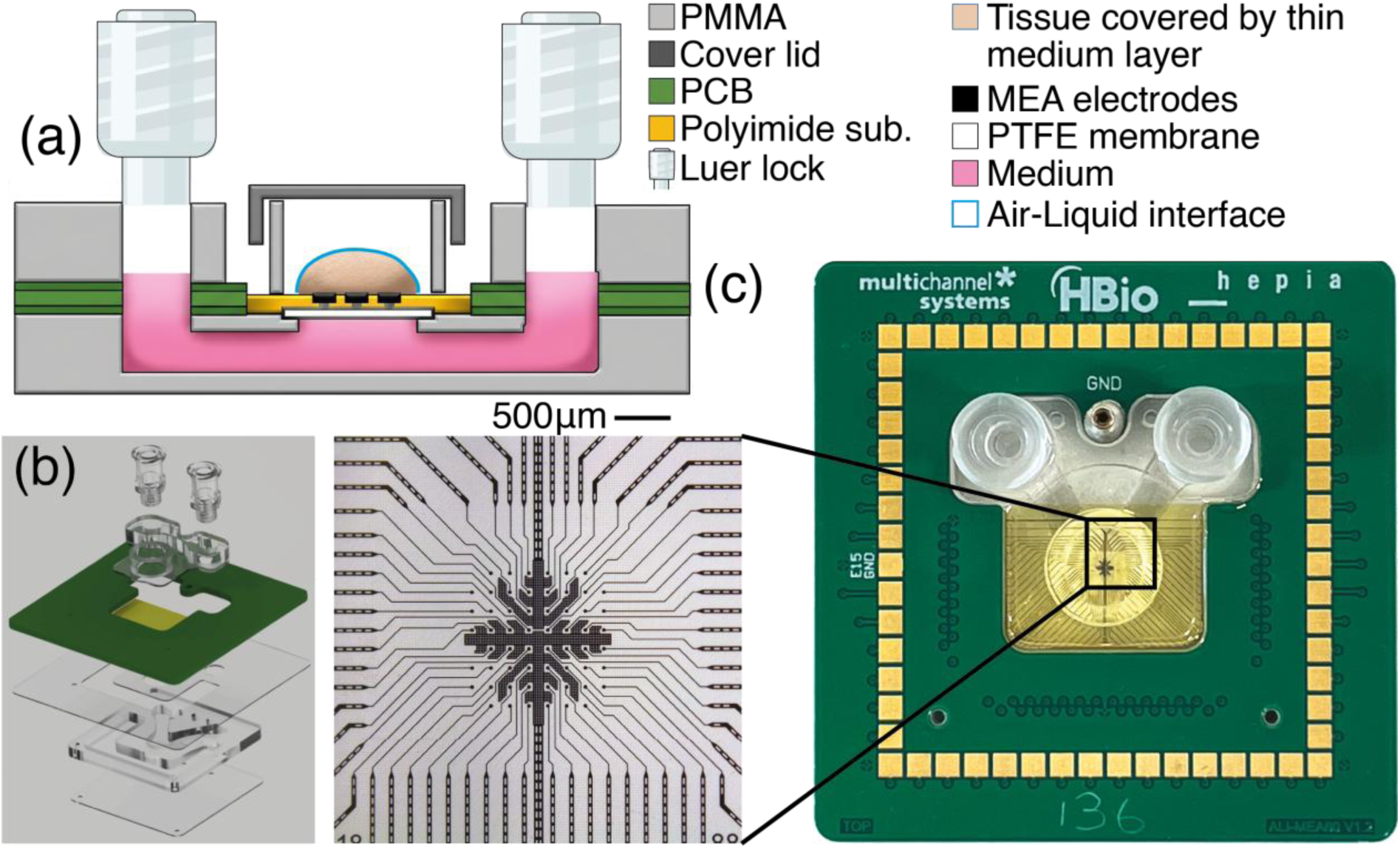
The ALI MEA platform: design, integration and electrical specifications. (a) Schematic cross-section of the assembled device. A 3D tissue (beige) is placed directly on the porous polyimide membrane (orange), with culture medium (pink) filling the basal chamber and a PTFE membrane (white) homogenizing the basal flow. The apical surface of the tissue is exposed to humidified air (air–liquid interface, blue outline). Medium is exchanged through the two Luer-lock ports. Materials are identified in the legend at the top right. (b) Exploded view of the device showing, from top to bottom, the PMMA cover lid with two Luer-lock ports, the porous polyimide thin-film MEA mounted on its PCB carrier, and the base housing forming the basal perfusion chamber. (c) Right: Assembled biochip (5×5cm) and close-up on brightfield micrograph of the active area, showing the 8 × 8 electrode grid (corner positions omitted) and the radial routing of the platinum tracks inside the polyimide membrane.

The recording substrate is an 8 µm-thick polyimide membrane perforated with through-holes of 7.5 µm diameter at 20 µm pitch, yielding an open porosity of approximately 10% across the active area. The membrane carries 59 recording/stimulation electrodes plus one reference electrode, arranged on an 8 × 8 grid (corner positions omitted) with 200 µm electrode pitch and 30 µm electrode diameter, defining a 1.4 × 1.4 mm^2^ active footprint matched to the typical dimensions of human neural organoids (Figure 1(c) - Left). The porous geometry ensures that culture medium delivered to the basal chamber (350 µL volume, accessible via two Luer-lock ports) can diffuse vertically through the membrane and reach the basal face of the tissue, while the apical surface remains exposed to humidified air. A hydrophilic PTFE membrane (0.45 µm pore size) sealing the basal chamber homogenises basal flow and prevents bubble formation during medium exchange.

Electrochemical performance was characterised across the production population (N= 41 chips, 59 electrodes per chip) in physiological saline at 1 kHz, after Pt-black electrodeposition (Table 1). Per-chip impedance midpoints, computed as the centre of the [min, max] range across the 59 electrodes of each chip, had a population median of 56 kΩ with an interquartile range of 42–82 kΩ, and a full population range of 24–107 kΩ. Per-chip phase midpoints were tightly distributed around a population median of 64° (IQR 58–68°, range 45–73°), consistent with the expected behaviour of Pt-black-coated microelectrodes in the kHz range. The reference electrode, fabricated from the same material with a larger geometric area, exhibited an impedance of 2.41 ± 0.52 kΩ (mean ± SD) across the N = 41 chips. This narrow distribution across a population of independently fabricated and post-processed chips supports the reproducibility of the manufacturing and electrodeposition processes.

**Table 1:** Summary specifications of the platform. Geometry and membrane parameters describe the device as fabricated. Electrochemical specifications are reported across the production population (N = 41 chips, 59 recording electrodes per chip), measured in physiological saline at 1 kHz after Pt-black electrodeposition; per-chip impedance and phase values are reported as the midpoint of the [min, max] range across the 59 electrodes of each chip, with the population summarized by its median and interquartile range (IQR). The reference electrode impedance is reported as mean ± SD across the N = 41 chips. RMS noise was measured on representative chips in physiological saline and in culture medium under standard acquisition conditions.

| Geometry & layout |  |
|---|---|
| Recording/stimulation electrodes | 59 |
| Reference electrode | 1 |
| Arrangement | 8 × 8 grid (corners omitted) |
| Electrode diameter | 30 *µ*m |
| Electrode pitch | 200 *µ*m |
| Active area footprint | 1.4 × 1.4 mm² |
| Membrane |  |
| Material | Polyimide |
| Thickness | 8 *µ*m |
| Through-hole diameter | 7.5 *µ*m |
| Through-hole pitch | 20 *µ*m |
| Open porosity | ~10 % |
| Electrochemistry (N = 41 chips, 59 electrodes per chip) |  |
| Electrode coating | Platinum black |
| Impedance at 1 kHz, per-chip midpoint | median 56 kΩ (IQR 42-82) |
| Impedance at 1 kHz, population range | 24-107 kΩ |
| Phase at 1 kHz, per-chip midpoint | median 64° (IQR 58-68) |
| Phase at 1 kHz, population range | 45-73° |
| Reference electrode impedance | 2.41 ± 0.52 kΩ (mean ± SD) |
| Fluidics |  |
| Apical configuration | Air-Liquid interface |
| Basal chamber access | Two Luer-lock ports |
| Basal chamber volume | 350 *µ*L |
| Acquisition compatibility |  |
| System | MEA2100-Mini (MCS) |
| Sampling rate | Whole available range of MCS system |
| Noise (RMS, saline / culture) | ±5 *µ*V in saline; 10-20*µ*V in medium |

Recording-grade noise performance was verified under standard acquisition conditions on representative chips. Root-mean-square noise was approximately ±5 µV in physiological saline and 10–20 µV when chips were filled with culture medium, the latter consistent with the higher conductivity and biological content of standard NDM. Both values are below the 25 µV ceiling commonly used to flag noisy channels in extracellular recordings and well within the dynamic range required for reliable detection of unitary events (typically 20–50 µV peak-to-peak) in 3D cultures.

Together, these geometric, electrochemical, and noise characteristics define a recording substrate that is (i) reproducibly fabricated across a population of 41 independent chips, (ii) electrochemically suited for extracellular recordings at the µV scale, and (iii) physically compatible with the maintenance of 3D tissues at the air-liquid interface. The remainder of this study assesses the functional consequences of this design for chronic recording, pharmacological responsiveness, and cross-tissue applicability.

### 3.2 Spike band and local field potential activity from 3D neural cultures

We first placed human neural spheroids on the ALI MEA active area and acquired extracellular activity to establish the platform’s recording capability under physiological conditions (Figure 2(a)). Spike-band filtering of the wideband signal (300–3000 Hz) revealed clear unitary events distributed across the array (Figure 2(b)). The spatial firing-rate map showed marked heterogeneity, with active sites clustered in regions of tissue contact and quiescent regions in areas not covered by tissue (Figure 2(c)). Mean firing rates across QC-good electrodes ranged from approximately 0.2 to >10 spikes·s^-1^, with the upper end of the colour scale clipped at 15 spikes·s^-1^ to preserve dynamic range. Channels rejected by QC (i.e., no detected spikes) are shown in white.

**Figure 2:**
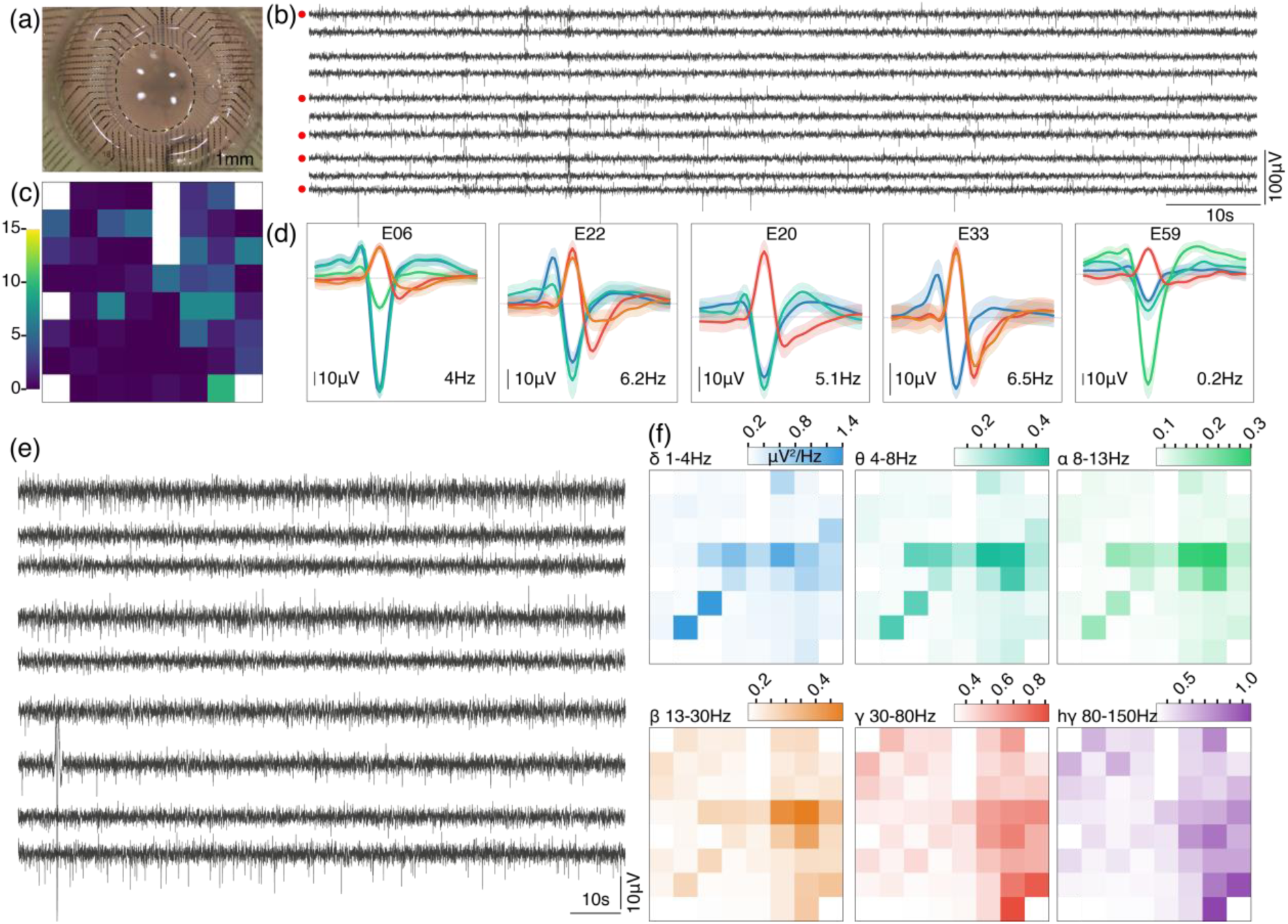
Extracellular recording of a neuronal culture on the ALI MEA60 platform. **(a)** Neural spheroids plated on the active area; scale bar 1 mm. **(b) Spike-**band traces (band-pass 300–3000 Hz) from 10 representative electrodes over a 2-min window; red dots mark the five electrodes shown in (d). (c) Mean firing rate per electrode on the 8×8 layout (scale 0–15 Hz); white cells = QC-rejected channels. (d) Sorted spike waveforms for five electrodes. Sorting: PCA (3 components) + GMM (BIC-optimal k, max 4 units per polarity); extraction window −1 / +1.5 ms around the absolute peak. Thick line = mean, shaded band = ±1sd. Blue/green = negative-polarity units, red/orange = positive-polarity units; per-electrode firing rate indicated bottom-right of each panel. (e) LFP/wideband traces (band-pass 1–200 Hz) from the same 10 electrodes as in (b). (f) Spatial maps of mean band power across the 8×8 array for six canonical bands: δ (1–4 Hz), θ (4–8Hz), α (8–13 Hz), β (13–30 Hz), γ (30–80 Hz), high-γ (80–150 Hz). Colour scales independent per band (μV²/Hz) to emphasize within-band spatial distribution.

Spike sorting on five representative electrodes resolved multiple putative units per recording site (Figure 2(d)). Both negative- and positive-polarity components were recovered, often co-existing on the same electrode, consistent with the presence of multiple cell bodies and neurites within the effective recording radius of the 30-µm Pt-black electrodes. Peak-to-peak waveform amplitudes were typically in the 40-80 µV range, well above the recording noise floor of the platform in culture medium (10-20 µV RMS), and per-electrode firing rates within the sorted units ranged from 0.2 to 6.5 Hz across the five examples shown.

Wideband filtering of the same recordings (1-200 Hz) yielded continuous low-frequency activity distinct from the bursty patterns observed in the spike band (Figure 2(e)). Decomposition into six canonical frequency bands: δ (1-4 Hz), θ (4-8 Hz), α (8-13 Hz), β (13-30 Hz), γ (30-80 Hz), and high-γ (80-150 Hz) showed that mean band power was spatially structured across the array, and that the spatial pattern differed between bands (Figure 2(f)). Lower-frequency bands (δ, θ) concentrated their power in localised regions of the array, while higher-frequency bands (γ, high-γ) showed broader, more distributed patterns. Each band is displayed on an independent colour scale to emphasise within-band spatial distribution rather than absolute cross-band magnitudes.

The ALI MEA at the ALI thus resolves both fast unitary events and slow population-level oscillations within a single recording session, giving a multi-scale read-out of network activity from one 3D culture.

### 3.3 Chronic recording stability across 60 days

Long-term stability is a prerequisite for using the ALI MEA60 in chronic pharmacological or developmental studies. We therefore monitored signal quality and channel-level network metrics on N = 7 chips over 60 days, comprising 121 recordings in total; two outlier recordings were excluded by a 3 × MAD intra-chip criterion applied independently to each metric.

Recording noise remained stable throughout the entire follow-up window. The across-chip median noise RMS hovered around 6-8 µV with no systematic drift (Figure 3(a)), and a linear mixed-effects (LME) model yielded a slope indistinguishable from zero (β = +0.004 µV/day, p = 0.87, chip identity as random intercept). Median noise remained below the 15 µV QC threshold at every time point, with only sporadic excursions on individual chips and recordings. This indicates that recording noise remained stable over the two-month culture period, with no evidence of progressive degradation in recording quality.

**Figure 3:**
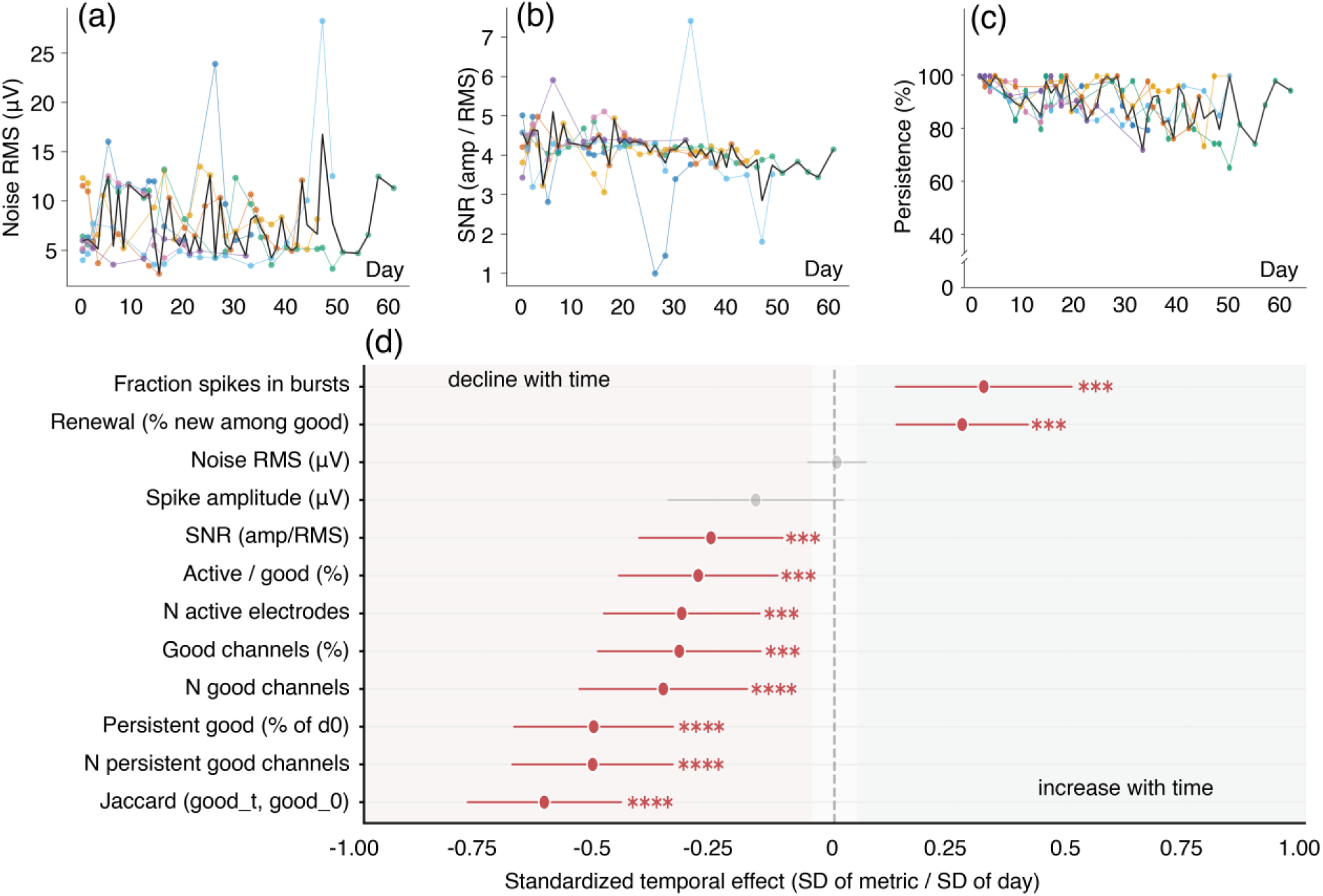
Long-term recording stability of the ALI MEA over 60 days (N = 7 chips, 121 recordings; 2 outlier recordings excluded by a robust 3·MAD intra-chip criterion). (a–d), coloured lines represent individual chips and the black line the across-chip median. (a) Noise RMS remains below the 15 μV quality threshold throughout the recording period (LME β = +0.004 μV/day, p = 0.87). (b) SNR shows a marginal decline (β = −0.013/day, p = 0.001) but median values remain at or above the SNR ≥ 4 threshold. (c) Persistence: the fraction of Day-0 good channels still QC-good at later time points stays above 75% (median ≈ 85%). (d) Standardized temporal effects (β × SD (day) / SD (metric)) for 12 representative metrics, ordered by effect magnitude. Negative values indicate metrics that decrease with time (left), positive values metrics that increase with time (right); the dashed line at zero indicates no temporal trend. Significance levels: *p < 0.05, **p < 0.01, ***p < 0.001, ****p < 0.0001. Pharmacological validation across three compound families

SNR defined as the ratio of mean detected spike amplitude to noise RMS, showed a small but statistically detectable decline over time (β = −0.013/day, p = 0.001; Figure 3(b)). Despite this trend, the across-chip median remained at or above the SNR ≥ 4 threshold throughout the 60-day window, and the absolute magnitude of the effect (∼0.8 SNR unit over the full follow-up) is well within the dynamic range needed for reliable spike detection. The decline reflects a modest reduction in spike amplitude rather than a rise of the noise floor, which remained stable (Figure 3(a)). Because RMS noise remained stable over the same window, this amplitude reduction is more consistent with tissue-side factors (*i.e.*, maturation-related reorganisation of active sites and evolving tissue–electrode coupling) than with degradation of the electrodes or their Pt-black coating.

Persistence, defined as the fraction of channels classified as QC-good at Day 0 that remained QC-good at later time points, stayed above 75 % throughout the recording window, with an across-chip median of approximately 85 % (Figure 3(c)). Channels that produced high-quality recordings on the first day therefore continued to do so two months later, indicating that channel-level functionality is preserved on the same physical electrodes over chronic culture.

To consolidate these per-metric observations into a single readout, we computed standardized temporal effects (β × SD (day) / SD (metric)) from per-metric LME models for 12 representative metrics spanning signal quality, channel yield, and network organisation (Figure 3(d)). Three patterns emerged. First, the two raw signal-quality metrics: noise RMS and mean spike amplitude showed no significant temporal trend, confirming the visual stability seen in Figure 3(a). Second, channel-yield metrics (N active electrodes, N good channels, Active/good %, Good channels %, SNR) showed a modest but consistent decline (|effect| ≈ 0.20–0.35 SD, all p < 0.001), reflecting a slow attrition of usable channels rather than a collapse of recording quality. Third, channel-identity-tracking metrics (Persistent good, N persistent good channels, Jaccard with Day 0) showed the largest declines (|effect| ≈ 0.45–0.60 SD, p < 0.0001), as expected when the *identity* of good channels is required to remain stable rather than merely their *number*. Two metrics increased over time: the fraction of spikes occurring in bursts and the renewal rate (the fraction of currently good channels that were not good at Day 0), both consistent with progressive network maturation and reorganisation of active sites across the array.

Overall, the ALI MEA60 supports a recording window of at least 60 days during which (i) noise floor and SNR remain compatible with reliable spike detection, (ii) the majority of channels recorded on Day 0 continue to deliver usable signals two months later, and (iii) longitudinal metrics evolve along interpretable axes — slow channel attrition combined with ongoing network maturation — rather than abrupt failure modes.

To demonstrate that the ALI MEA60 captures network-level responses to bioactive compounds, we applied a sequential pharmacological protocol to each chip (N = 7), targeting three distinct synaptic mechanisms: gamma-aminobutyric acid A (GABA_A_) receptor blockade (gabazine, 5 µM), voltage-gated K^+^ channel blockade (4-aminopyridine [4-AP], 100 µM), and combined α-amino-3-hydroxy-5-methyl-4-isoxazolepropionic acid (AMPA) / *N*-methyl-D-aspartate (NMDA) glutamate receptor blockade (6-cyano-7-nitroquinoxaline-2,3-dione (CNQX) 20 µM + D-2-Amino-5-phosphonovaleric acid (D-AP5) 50 µM). Each compound was preceded and followed by Baseline NDM recordings, with wash periods between conditions (Figure 4(a)). Four network metrics were quantified at the recording level: burst rate (bursts·min^-1^), proportion of spikes occurring within bursts, mean firing rate (spikes·s^-^¹), and the mean Spike Time Tiling Coefficient (STTC) among the top 10 % most correlated electrode pairs as a proxy for functional connectivity within the most coactive subnetwork.

**Figure 4:**
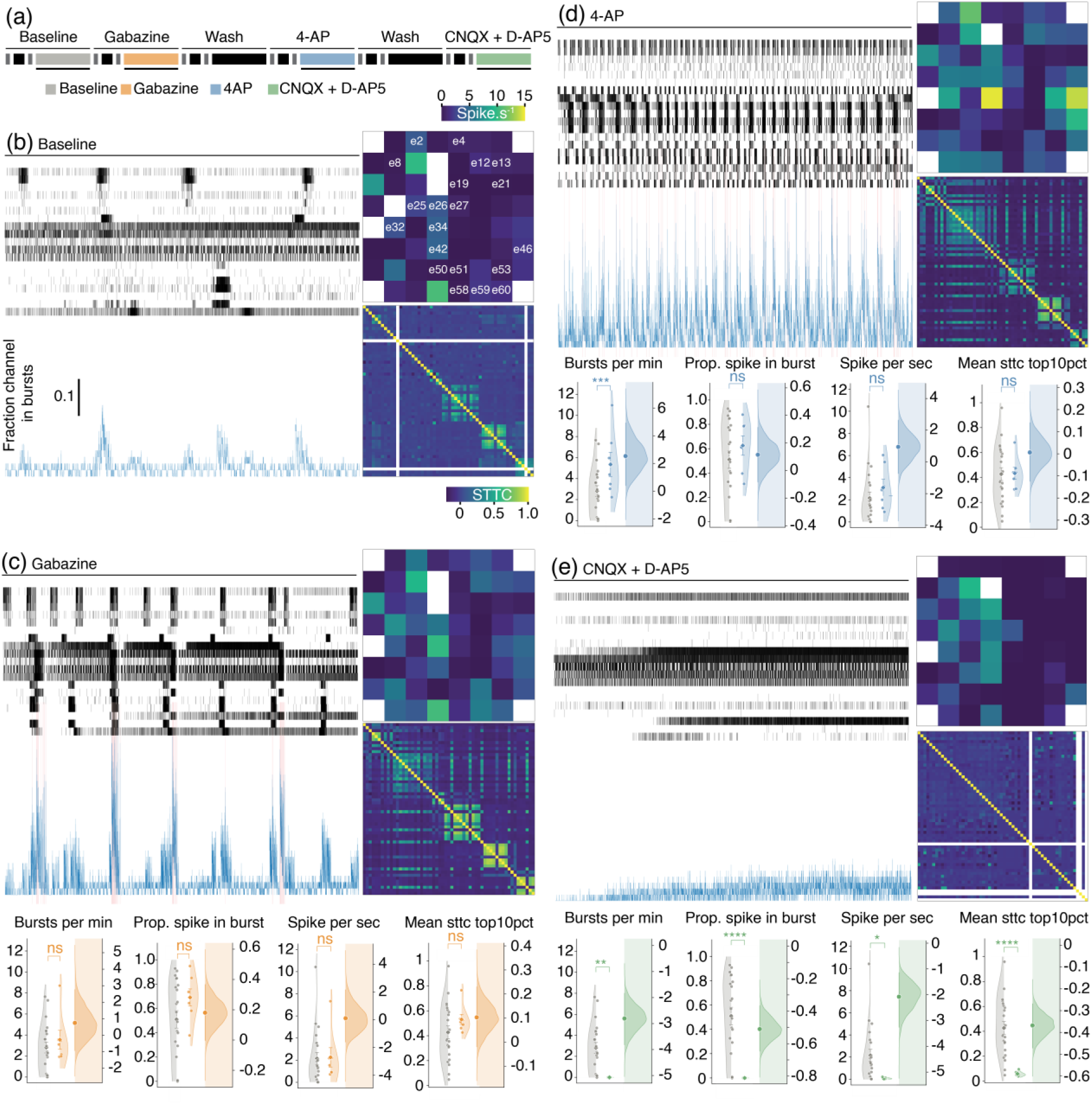
MEA chip pharmacological validation across three compound families. (a) The condition timeline shows the sequential pharmacological protocol applied to each chip: Baseline NDM → Gabazine → Wash → 4-AP → Wash → CNQX + D-AP5. Colours indicate the pharmacological family (grey: Baseline, orange: Gabazine, blue: 4-AP, green: CNQX + D-AP5). (b-e) Representative recordings from a single chip under four conditions: Baseline NDM (b), Gabazine (c), 4-AP (d), and CNQX + D-AP5 (e). For each condition, the upper panel shows the spike raster (one row per electrode; the 20 electrodes displayed in white labels correspond to those represented in the firing-rate heatmap). (b) raster plot with the corresponding population rate trace below (blue, smoothed Gaussian, bin = 10 ms). The insert heatmap (upper right) displays the spike frequency per second across all the electrodes (colour scale 0–15 spikes·s^-1^), and the lower panel: Spike Time Tiling Coefficient (STTC) matrix across all good-quality electrode pairs shows the spatial organization of pairwise functional connectivity. (lower panels). Quantification of four network metrics for each pharmacological family, shown as paired scatter + half-violin plots (left axis) and bootstrapped estimation distributions (right axis, shaded). Each dot represents the recording-level median across good-quality electrodes for one chip (N = 7). The diamond symbol indicates the mean ± SEM. The shaded estimation distribution (right of each panel pair) shows the bootstrap distribution of the mean difference (drug − Baseline NDM, 5,000 iterations) with the 95% confidence interval indicated by the vertical bar and the observed difference by the filled circle. Metrics show burst rate (bursts per minute), proportion of spikes occurring within bursts (prop. spike in burst), firing rate (spikes per second), and mean STTC among the top 10% most correlated electrode pairs (mean STTC top10pct). Statistical annotations indicate the result of linear mixed model (LME) comparisons at the electrode level (drug versus Baseline NDM; chip identity as random intercept, recording as variance component; Holm correction within model): ns p ≥ 0.05, * p < 0.05, ** p < 0.01, *** p < 0.001, **** p < 0.0001.

#### Baseline

Under control conditions, recordings showed sparse spontaneous spiking distributed across the array, organised into infrequent and weakly synchronised network events. The population rate trace exhibited low-amplitude fluctuations with occasional discrete bursts (Figure 4(b)), the firing-rate heatmap revealed a small number of moderately active sites, and the STTC matrix showed limited pairwise synchrony confined to a small cluster of electrodes. These baseline characteristics were consistent across the seven chips and provided the reference condition against which all subsequent drug effects were compared.

#### Gabazine

Disinhibition by GABA_A_ receptor blockade reorganised network activity into clearly synchronised bursts. The raster showed coordinated population events repeating throughout the recording, accompanied by sharp peaks in the population rate trace (Figure 4(c), pink shading). The firing-rate heatmap retained a similar spatial distribution to baseline, while the STTC matrix exhibited visibly stronger and more widespread off-diagonal correlations, indicating that disinhibition recruited a larger fraction of electrodes into synchronous events. At the population level, however, none of the four LME-tested metrics reached statistical significance versus baseline (all p ≥ 0.05; Figure 4(c), lower panels). This is consistent with gabazine acting primarily on temporal organisation, converting tonic firing into synchronised bursts, rather than on the amount of activity quantified by spike or burst rates averaged over the recording.

#### 4-AP

K+ channel blockade by 4-AP produced a pronounced increase in network excitability. The raster showed dense, near-continuous spiking across most electrodes, with the population rate elevated throughout the recording (Figure 4(d)). Burst rate increased significantly relative to baseline (p < 0.001, LME), while firing rate, proportion of spikes in bursts, and mean STTC top10 % did not reach significance at the electrode level (all p ≥ 0.05). The bootstrap distribution of the mean difference for burst rate was clearly displaced from zero, confirming the robustness of the effect.

#### CNQX + D-AP5

Combined AMPA/NMDA glutamate receptor blockade produced the expected silencing of glutamatergic network activity. The raster collapsed to sparse, asynchronous activity on a small subset of electrodes (Figure 4(e)), and the population rate flattened to near-zero. Four network metrics decreased significantly *versus* baseline: burst rate (p < 0.01), proportion of spikes in bursts (p < 0.0001), firing rate (p < 0.05), and mean STTC top10 % (p < 0.0001), with bootstrap distributions of the mean difference shifted well into the negative range. The residual STTC structure visible in the matrix likely reflects shared input or remaining non-glutamatergic activity rather than glutamate-mediated functional connectivity.

#### Statistical consistency

All effects reaching significance under the electrode-level LME analysis (chip identity as random intercept, recording as variance component, Holm-corrected within each model) were confirmed in direction by recording-level Friedman tests, supporting the robustness of the observed responses to the choice of statistical framework.

The ALI MEA60 is therefore able to capture the canonical signatures of three distinct pharmacological perturbations: disinhibition-driven network synchronisation (gabazine), broad excitability enhancement (4-AP), and glutamatergic silencing (CNQX + D-AP5). The platform therefore captures both quantitative changes in firing and burst rates and qualitative reorganisations of network synchrony, providing a foundation for its use as a functional readout in chronic drug-screening applications.

### 3.4 Cross-tissue extension of the ALI MEA60 Platform

A practical consequence of recording at the ALI on a porous polyimide membrane is that the same device geometry accommodates tissues of widely different origin, thickness, and physiology, without modifying the protocol beyond the choice of culture medium and acquisition window. To demonstrate this versatility, we used the ALI MEA60 with three preparations that cover the representative use cases of extracellular electrophysiology: human brain slices, human brain organoids (3D neural samples) and human cardiac organoids (a 3D non-neural preparation with intrinsic beating activity).

#### Human brain slices

Spontaneous spiking activity was recorded from human brain slices placed on the ALI MEA60. The spatial firing-rate map across the 8 × 8 array showed a focal hotspot of activity reaching ∼5 spikes·s^-^¹ on a single electrode, surrounded by a broader region of moderately active sites in the 1–3 Hz range (Figure 5(a), left); the corresponding map of the number of putative units per electrode showed that isolated units were recovered across a spatially distributed subset of sites (Figure 5(a), right). Spike sorting on four representative electrodes recovered well-isolated unitary events with peak-to-peak amplitudes of 20–40 µV and stereotyped waveform shapes including both predominantly negative and biphasic profiles (Figure 5(c)), indicating that the platform resolves individual units in human tissue with signal quality comparable to that obtained on neural spheroid cultures. Over a 30-min recording, the raster across all 60 channels revealed sustained spiking activity on a stable subset of electrodes coexisting with sparser, spatially distributed events on the remainder (Figure 5(b)). Beyond spiking, band-limited power maps computed across the array in six canonical frequency bands (δ 1–4, θ 4–8, α 8–13, β 13–30, γ 30–80 and high-γ 80– 150 Hz) revealed spatially structured, band-specific power distributions (Figure 5(d)), demonstrating that the preparation supports both single-unit-level and population-level (field) read-outs on the same device.

**Figure 5:**
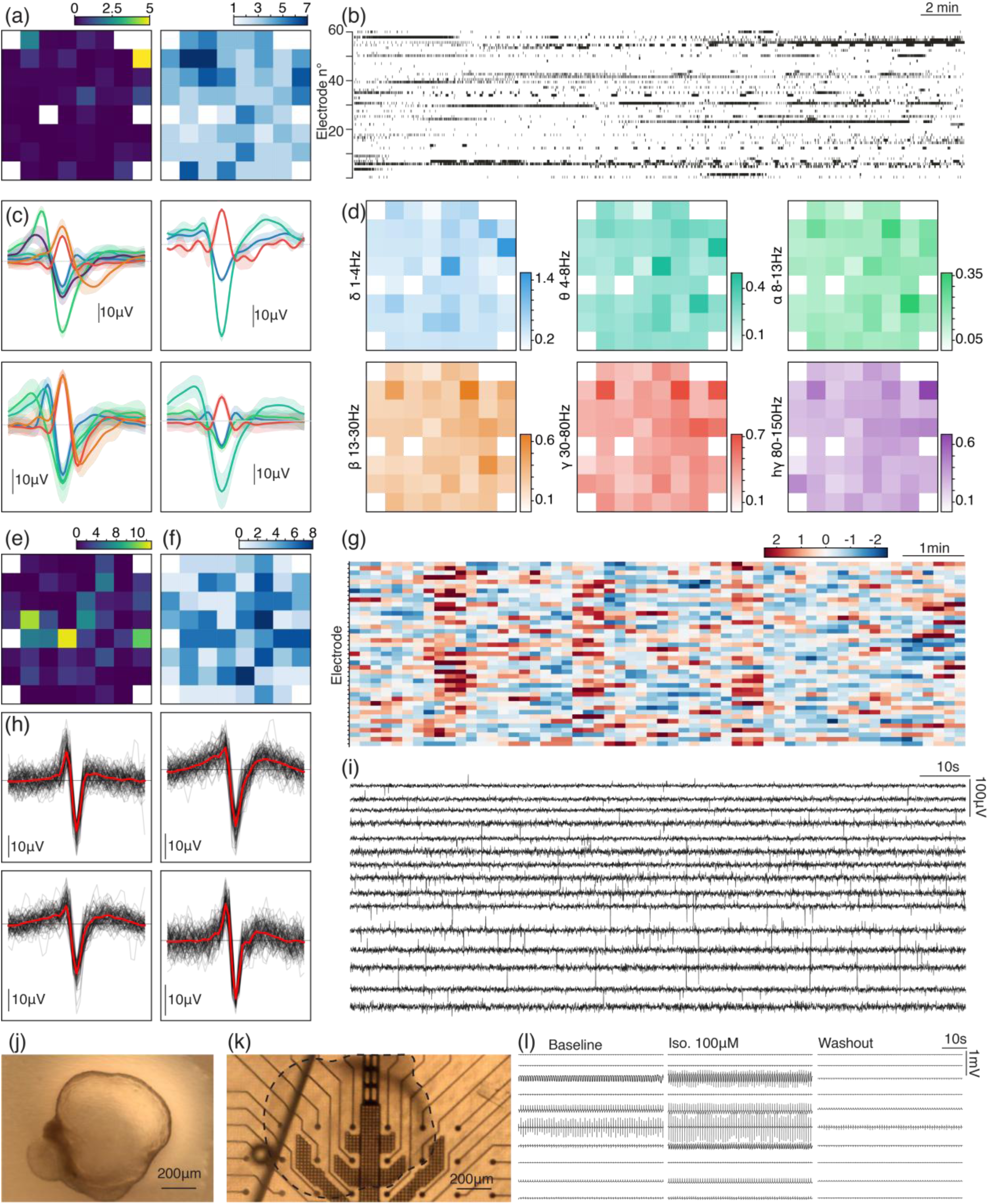
Cross-tissue extension of the ALI MEA60 platform. (a–d) Ex vivo human brain slices. (a) Spatial firing-rate map across the 8×8 array (left; scale 0–5 Hz) and number of putative units per electrode (right: scale 1–7). (b) Raster of detected spikes across the 60 channels over a 30-min recording. (c) Mean spike waveforms (coloured lines) with shaded ±sd for the sorted units of four representative electrodes; scale bar 10 μV. (d) Spatial distribution of band-limited power across the array in six canonical frequency bands (δ 1–4 Hz, θ 4–8 Hz, α 8–13 Hz, β 13–30 Hz, γ 30–80 Hz, high-γ 80–150 Hz); colour scales independent per band. (e–i) Human brain organoid. (e) Spatial firing-rate map (8×8; scale 0–10 Hz). (f) Number of putative units per electrode (scale 0–8). (g) Firing-rate stability across the recording, expressed as a per-electrode Z-score over time (scale ±2; 1-min bins). (h) Mean spike waveforms (red) and individual events (grey, ∼25 events superimposed) for four representative electrodes; time window 2 ms. (i) Spike-band traces (band-pass 300–3000 Hz) from 15 active electrodes; scale bars 10 s (horizontal), 100 μV (vertical). (j) Cardiac organoid prior to plating. (k) Same organoid (dashed outline) placed on the porous polyimide membrane over the array. (l) Field-potential traces (band-pass 0.5–100 Hz for display) from the six electrodes with highest beating activity, under three sequential conditions: Baseline → Isoproterenol (100 μM, 10 min wash-in) → Washout (10 min after removal). Traces are simultaneous across conditions and recorded from the same electrodes on the same chip.

#### Human brain organoids

To extend the platform to stem-cell-derived 3D neural tissue, we recorded spontaneous activity from single human brain organoids placed at the ALI on the ALI MEA60. The spatial firing-rate map across the array revealed a spatially structured activity pattern, with a subset of electrodes beneath the organoid reaching firing rates of up to ∼10 Hz and surrounding sites showing lower-rate activity (Figure 5(e)). Spike sorting across active electrodes identified up to 8 putative units per electrode (Figure 5(f)), indicating that several single units can be resolved simultaneously from the same organoid. Firing-rate stability, expressed as a per-electrode Z-score in 1-min bins across the recording, showed stable activity over the recording window (Figure 5(g)). Waveform extraction on four representative electrodes recovered well-isolated events with stereotyped, predominantly biphasic spike shapes (Figure 5(h); peak-to-peak amplitudes ∼20–40 µV, 2-ms window). Spike-band traces (300–3000 Hz band-pass) from 15 active electrodes confirmed extracellular spikes rising clearly above the noise floor across the array (Figure 5(i)), demonstrating that the ALI MEA60 resolves single- and multi-unit activity from human brain organoids under the same acquisition conditions used for the other preparations.

#### Human cardiac organoids

To test whether the platform extends to non-neural 3D tissues with intrinsic rhythmic contractile activity, we placed single human cardiac organoids on the polyimide membrane at the ALI (Figure 5(j,k)). Field-potential recordings from the six electrodes with the highest beating activity exhibited regular, large-amplitude deflections at approximately 1 Hz under baseline conditions (Figure 5(l), left; band-pass 0.5–100 Hz for display). Application of the β-adrenergic agonist isoproterenol (100 µM, 10-min wash-in) produced a visible chronotropic acceleration on the same electrodes, consistent with the expected positive control for β-adrenergic responsiveness in cardiac tissue (Figure 5(l), centre). The effect reversed after a 10-min washout to partial recovery of baseline rhythm (Figure 5(l), right). These traces were recorded simultaneously across the six electrodes from a single chip, demonstrating that the porous polyimide membrane transduces field-potential signals from cardiac tissue with sufficient amplitude and bandwidth to resolve pharmacologically induced rate changes.

#### Cross-preparation summary

Across the three preparations, the ALI MEA60 captured the principal modes of extracellular signal expected from each tissue: multi-unit spiking and network-burst dynamics from human brain slices, single- and multi-unit activity from human brain organoids, and rhythmic field potentials from cardiac organoids. No modification was made to the device, its electrode configuration, the basal perfusion chamber, or the acquisition chain between preparations; only the culture medium and recording duration were adapted to each tissue type. This versatility, combined with the chronic stability and the pharmacological responsiveness, supports the use of the ALI MEA60 as a generic platform for functional electrophysiology on 3D and *ex vivo* tissues across neural and non-neural contexts.

## 4 DISCUSSION

We have introduced and characterised the ALI MEA60, a porous polyimide substrate-based thin-film microelectrode array that maintains 3D neural and non-neural tissue at the ALI while remaining compatible with the commercially available MEA2100-Mini data acquisition system. Its electrochemical profile was reproducible across N = 41 chips: chronic recordings on human neural spheroids remained usable for at least 60 days, the platform reproduced the canonical signatures of gabazine, 4-AP and CNQX/D-AP5 and the same device recorded from human brain organoids, slices and human cardiac organoids.

Its distinguishing feature is to combine, on a single device, three capabilities that conventional MEAs typically trade off against one another: ALI culture compatible with 3D tissue maintenance^18,19^, chronic recording stability over weeks to months, and applicability across neural and non-neural tissue.

### 4.1 The air–liquid interface as an enabler of 3D and *ex vivo* electrophysiology

The defining design choice of the ALI MEA60 is that the recording substrate is a thin (8 µm), perforated polyimide membrane rather than a glass slide or a closed-bottom MEA dish. This design carries three coupled consequences. First, at the ALI the tissue is not submerged but covered only by a thin film of medium maintained by capillarity, which keeps the preparation hydrated while allowing direct gas exchange between this film and the humidified incubator atmosphere^4,17^. This configuration supports far more efficient tissue oxygenation than diffusion through a bulk medium column, thereby reducing the diffusion limitations associated with central necrosis in long-term submerged 3D cultures^18,25–27^. Second, culture medium delivered to the basal chamber reaches the basal face of the tissue through the membrane pores, providing a continuous nutrient and gas exchange path that does not rely on apical diffusion. Third, tissues adhere directly to the porous polyimide surface: no coating or adhesion promoter was required for stable attachment across any of the preparations tested, simplifying preparation and removing a common source of batch-to-batch variability. The combination is, to our knowledge, what allows the same platform to sustain neural spheroids for >60 days^16–18^ and to accommodate organotypic human cortical slices, human brain organoids and cardiac organoids without modification of the device or its perfusion chain.

### 4.2 Stability over 60 days: a tractable failure mode

The longitudinal characterisation provides what we believe is the most quantitatively defensible part of this work. Rather than reporting a binary “stable up to day X” claim, we tracked 12 metrics across the cohort and quantified each as a standardised temporal effect with Holm–Bonferroni correction control. The pattern that emerged is informative: noise RMS was temporally flat and mean spike amplitude showed only a slight, non-significant decline, while metrics capturing channel-level identity (Jaccard with Day 0, Persistent good, N persistent good channels) declined more strongly than metrics capturing channel-level counts (N good channels, N active electrodes). In other words, the platform loses channel identity faster than it loses channel quantity — i.e., the set of good channels at Day 60 partially overlaps with, but is not identical to, the set at Day 0.

This is a tractable failure mode. For longitudinal studies that rely on tracking the same units over time (developmental electrophysiology, chronic pharmacological titration), it sets a quantitative ceiling on how long the same channels can be relied upon, with a soft floor at the level of population-level metrics that remain stable. For applications that only require a sufficient yield of good channels at each time point (drug screening, network-level characterisation), the ceiling is correspondingly higher. The complementary rise of the “Renewal” metric (fraction of currently good channels that were not good at Day 0) further suggests that the cultures themselves continue to mature and reorganise across the recording window, contributing to the observed turnover of active sites. Together with the stable noise floor, this points to biological reorganisation — different electrodes sampling activity on different days — rather than electrode-level drift, as the main driver of good-channel turnover.

### 4.3 Pharmacological responsiveness and the limits of rate-based metrics

The pharmacological validation reproduced the expected qualitative effects of three reference compound families, but with a notable asymmetry in statistical detectability. CNQX + D-AP5 produced highly significant decreases across all four tested metrics, 4-AP produced a significant increase in burst rate alone, and gabazine — despite a visually unambiguous reorganisation of network activity into synchronised bursts — did not reach significance on any of the four metrics tested. These directions are consistent with prior MEA studies on human iPSC-derived and dissociated neuronal networks, in which glutamatergic blockade abolishes network bursting while GABA_A blockade reorganises activity toward more synchronised bursting rather than uniformly scaling overall firing^28,29^. This asymmetry therefore deserves interpretation rather than dismissal.

The four metrics we quantified (burst rate, fraction of spikes in bursts, firing rate, mean STTC of the top 10 % most correlated pairs) are rate-based or integrated summaries of network activity. They are well-suited to capture changes in the amount of activity (CNQX abolishes spiking; 4-AP increases burst frequency) but less suited to capture changes in the temporal organisation of activity at a constant total spike count (i.e. precisely the regime suggested by the gabazine recordings presented here). This is a known limitation of rate-centric read-outs: in multi-parameter analyses of MEA drug responses, firing-rate variables contribute little to the discriminative signal relative to burst-structure and connectivity features^30,31^. Independent recording-level Friedman tests confirmed the direction of all LME-significant effects, ruling out a statistical artefact, but they cannot rescue an effect that the chosen summary does not encode. Demonstrating sensitivity to gabazine-like reorganisations will therefore require complementary metrics targeting burst structure (duration, inter-burst interval, spikes per burst), network-burst propagation and recruitment, and higher-order synchrony beyond top-pair STTC^32,33^. Building these into a routine analysis pipeline is a natural extension of the present work.

### 4.4 Cross-tissue extension and applicability

The cross-preparation experiments were intentionally limited in scope: a single representative chip per tissue type, demonstrating that the device records the expected mode of activity from each preparation. They are sufficient to establish feasibility but not to characterise tissue-specific performance, and we do not present them as such. Their interest lies in showing that the same device geometry, fabrication chain, and acquisition protocol accommodate preparations that, in conventional electrophysiology, would each typically require a dedicated platform: a planar MEA for dissociated culture, a MEA insert for *ex vivo* explants, and a separate substrate for cardiac organoid field potentials. For laboratories that work across more than one tissue type, the operational gains of a single-platform workflow are non-trivial.

Two observations from the cardiac experiment warrant cautious interpretation. First, the isoproterenol response was demonstrated on a single cardiac organoid, providing proof of pharmacological responsiveness rather than quantitative pharmacological characterization. Previous cardiac organoid MEA studies supporting stronger pharmacological conclusions have generally relied on larger experimental cohorts and/or dose–response protocols rather than a single exposure condition. For example, increased beating rates following isoproterenol stimulation have been reported using repeated samples or multiple organoids, highlighting the importance of broader sampling for quantitative assessment^12,14,34,35^.

Second, the post-washout recordings showed a marked reduction in spontaneous activity rather than a complete return to baseline rhythm. This observation should not be interpreted as evidence of irreversible functional alteration, as β-adrenergic responses to isoproterenol are known to diminish during agonist exposure through receptor desensitization and related regulatory processes, while recovery following agonist removal is reported to be variable and highly dependent on experimental conditions^36–38^. Consequently, the reduced post-washout activity observed here is compatible with an insufficient washout interval, partial β-adrenergic receptor desensitization, or other tissue-level factors, but the present data does not allow discrimination between these possibilities. We therefore report this finding as an observational result rather than as evidence for complete pharmacological reversibility.

### 4.5 Limitations and future directions

Several limitations of the present work are worth stating explicitly. (i) The longitudinal cohort consists of N = 7 chips from a single batch of neural spheroids; chip-to-chip and batch-to-batch variability across independent productions would refine the stability estimates reported here. (ii) The pharmacological validation was performed with single concentrations chosen as positive controls; dose–response characterisation will be required to use the platform for quantitative pharmacology. (iii) Cross-tissue extension was demonstrated on representative samples from each tissue type and does not constitute a systematic comparison with dedicated platforms for each preparation. Establishing the generality of the approach through extensive validation across additional 3D excitable tissues (e.g., spinal cord slices, retinal explants, or neuromuscular co-cultures) would require a substantially broader study and was intentionally beyond the scope of the present work.

Beyond addressing these limitations, three extensions appear particularly promising. First, integrating closed-loop stimulation with the existing recording capability would enable causal interrogation of network states in chronically maintained 3D cultures — a capability for which the ALI configuration is well-suited, since electrodes deliver charge into a tissue not isolated by a column of bulk medium. Second, coupling the platform with optical imaging through the porous polyimide membrane could combine simultaneous calcium or voltage imaging with extracellular electrophysiology, providing a multimodal read-out from the same 3D preparation. Third, enlarging the active recording area would better match spatially extended preparations such as *ex vivo* brain slices, whose footprint exceeds that of spheroids and organoids and is only partially covered by the current 8 × 8 array.

### 4.6 Conclusion

The ALI MEA60 occupies a previously underserved region of the design space for extracellular electrophysiology: 3D tissue compatibility, chronic recording stability, and cross-tissue versatility, on a single device. The chronic characterisation framework introduced here provides future users and developers with a quantitative yardstick for reporting device performance and comparing successive iterations.

## DATA AND CODE AVAILABILITY

The recording-level master dataset and the Python analysis pipeline used in this study are available from the corresponding author upon reasonable request.

## AUTHOR CONTRIBUTIONS

FFR: Conceptualization, Methodology, Software, Validation, Formal analysis, Investigation, Data Curation, Writing – Original, Writing – Review & Editing, Visualization | LST: Conceptualization, Methodology, Investigation, Writing – Review & Editing | SSI, MOH, LNI: Methodology, Writing – Review & Editing | ARO: Conceptualization, Methodology, Resources, Supervision, Project administration, funding acquisition, Writing – Review & Editing | TRI, MPU, MJA, LSC, AAM, LMO, ADC, MGU: Methodology, Review & Editing | JAC, JEM, SSC: Writing – Review & Editing

## ETHICS STATEMENT

### Human pluripotent stem cell lines

Human neural spheroids were generated from the commercially available human iPSC-derived neural stem cell line NSChiPS (Thermo Fisher Scientific). Human cardiac organoids were generated from the human embryonic stem cell line H9 (WA09; WiCell). Work with these established, commercially available human pluripotent stem-cell lines was carried out under Haute École Spécialisée de Suisse occidentale (HES-SO) approval and did not require additional review by an ethics committee.

### Human brain slices

*Ex vivo* human brain slices were obtained from patients undergoing resective surgery for pharmacoresistant epilepsy at Baggiovara Civil Hospital, with written informed consent from all donors, under a protocol approved by the local ethics committee (Comitato Etico Area Vasta Emilia Nord, prot. AOU 0007590/24). All procedures were performed in accordance with the Declaration of Helsinki and the Swiss Federal Act on Research involving Human Beings (Human Research Act, HRA).

## ACKNOWLEDGEMENTS

We are deeply grateful to the patients for generously donating tissue for research. We thank all participating surgeons and neurologists, particularly Dr. Giacomo Pavesi, Dr. Elisa Moriconi, Dr. Anna Vaudano and Dr. Stefano Meletti, for making human tissue neuroscience possible, and Prof. Jonathan Mapelli for helpful discussions.

## REFERENCES

1. Yang S, Hu H, Kung H, et al. Organoids: The current status and biomedical applications. MedComm. 2023;4(3):e274. doi:10.1002/mco2.274

2. Lampart FL, Iber D, Doumpas N. Organoids in high-throughput and high-content screenings. Front Chem Eng. 2023;5. doi:10.3389/fceng.2023.1120348

3. Fontanelli L, Nisini N, Pirola S, Recchia FA. Neuromuscular and cardiac organoids and assembloids: Advanced platforms for drug testing. Pharmacol Ther. 2025;272:108876. doi:10.1016/j.pharmthera.2025.108876

4. Tang J, Zheng H, Feng Y, Zeng J, Ma S. Attachment culture of cortical organoids at the microwell air-liquid interface. STAR Protoc. 2023;4(3):102502. doi:10.1016/j.xpro.2023.102502

5. Kopic I, Peng H, Schmidt S, et al. Inkjet-Printed 3D Sensor Arrays with FIB-Induced Electrode Refinement for Low-Noise Amperometric Recordings in hiPSC-Derived Brain Organoids. ACS Sens. 2025;10(9):6426–6435. doi:10.1021/acssensors.4c03740

6. Abu Shihada J, Jung M, Decke S, et al. Highly Customizable 3D Microelectrode Arrays for In Vitro and In Vivo Neuronal Tissue Recordings. Adv Sci. 2024;11(13):2305944. doi:10.1002/advs.202305944

7. Stoppini L, Buchs PA, Muller D. A simple method for organotypic cultures of nervous tissue. J Neurosci Methods. 1991;37(2):173–182. doi:10.1016/0165-0270(91)90128-m

8. Kim D, Ryu H. Recent Advances in Microelectrode Array Interfaces for Organoids. Biomimetics. 2026;11(2):142. doi:10.3390/biomimetics11020142

9. Mapelli L, Domenico DD, Sciacca G, et al. Enhanced electrophysiological recordings in acute brain slices, spheroids, and organoids using 3D high-density multielectrode arrays. PLOS ONE. 2025;20(9):e0328903. doi:10.1371/journal.pone.0328903

10. Trujillo CA, Gao R, Negraes PD, et al. Complex Oscillatory Waves Emerging from Cortical Organoids Model Early Human Brain Network Development. Cell Stem Cell. 2019;25(4):558–569.e7. doi:10.1016/j.stem.2019.08.002

11. Huang Q, Tang B, Romero JC, et al. Shell microelectrode arrays (MEAs) for brain organoids. Sci Adv. 2022;8(33):eabq5031. doi:10.1126/sciadv.abq5031

12. Kim K, Lee Y, Jung KB, et al. Highly Stretchable 3D Microelectrode Array for Noninvasive Functional Evaluation of Cardiac Spheroids and Midbrain Organoids. Adv Mater. 2025;37(6):e2412953. doi:10.1002/adma.202412953

13. Yuan Q, Qin C, Xu D, et al. PEDOT: PSS-Modified Organic Flexible and Implantable Microelectrode for Internal Bi-Directional Electrophysiology of Three-Dimensional Cardiomyocyte Spheroid. ACS Sens. 2025;10(1):460–469. doi:10.1021/acssensors.4c02755

14. Choi SJ, Liu Z, Yang F, et al. 3D Spatiotemporal Electrophysiology of Cardiac Organoids Using Shell Microelectrode Arrays. Adv Mater. 2026;38(2):e06793. doi:10.1002/adma.202506793

15. Bounik R, Landolt AE, Lee J, et al. Seamless integration of CMOS microsensors into open microfluidic systems. Lab Chip. 2025;25(9):2205–2221. doi:10.1039/d4lc01000k

16. Ozaki T, Ohta N, Ma J, Hirano M. 360° size-adjustable microelectrode array system for electrophysiological monitoring of cerebral organoids. Front Bioeng Biotechnol. 2025;13. doi:10.3389/fbioe.2025.1596009

17. Heuschkel MO, Stoppini L, Schoenecker S, et al. Air-liquid interface micro-electrode arrays - well-adapted tools for long-term in vitro monitoring of 3D neural tissues. Curr Dir Biomed Eng. 2025;11(1):266–269. doi:10.1515/cdbme-2025-0168

18. Stoppini L, Heuschkel MO, Loussert-Fonta C, Gomez Baisac L, Roux A. Versatile micro-electrode array to monitor human iPSC derived 3D neural tissues at air-liquid interface. Front Cell Neurosci. 2024;18. doi:10.3389/fncel.2024.1389580

19. Heuschkel MO, Stoppini L, Sivakumar S, Meents JE, Schoenecker S, Roux A. Porous 60-channel air-liquid interface micro-electrode arrays for long-term 3D neural tissues experimentation. 1cth Meet Neural Electrogenic Cell Interfacing. Published online 2025. Accessed July 15, 2026. https://arodes.hes-so.ch/record/16289

20. Govindan S, Batti L, Osterop SF, Stoppini L, Roux A. Mass Generation, Neuron Labeling, and 3D Imaging of Minibrains. Front Bioeng Biotechnol. 2021;8. doi:10.3389/fbioe.2020.582650

21. Hofbauer P, Jahnel SM, Papai N, et al. Cardioids reveal self-organizing principles of human cardiogenesis. Cell. 2021;184(12):3299–3317.e22. doi:10.1016/j.cell.2021.04.034

22. Yoon SJ, Elahi LS, Pașca AM, et al. Reliability of human 3D cortical organoid generation. Nat Methods. 2019;16(1):75–78. doi:10.1038/s41592-018-0255-0

23. Garcia S, Sprenger J, Holtzman T, Buccino AP. ProbeInterface: A Unified Framework for Probe Handling in Extracellular Electrophysiology. Front Neuroinformatics. 2022;16. doi:10.3389/fninf.2022.823056

24. Cutts CS, Eglen SJ. Detecting Pairwise Correlations in Spike Trains: An Objective Comparison of Methods and Application to the Study of Retinal Waves. J Neurosci. 2014;34(43):14288–14303. doi:10.1523/JNEUROSCI.2767-14.2014

25. Vijakumaran U, Anjum A, Devaraju P, Abd Razak NH, Daud MF, Mohd Idris I. Air–liquid interface–centered oxygen engineering in human brain organoids: intact, sliced, and microfluidic extensions. Front Cell Dev Biol. 14:1809928. doi:10.3389/fcell.2026.1809928

26. Cañizares Luna M, Mars M, Jakobs CE, et al. Protocol for generating and using human iPSC-derived microglia-containing air-liquid-interface cortical organoid cultures. STAR Protoc. 2025;6(3):103915. doi:10.1016/j.xpro.2025.103915

27. El Harane S, Nazari B, El Harane N, et al. Generation of Individualized, Standardized, and Electrically Synchronized Human Midbrain Organoids. Cells. 2025;14(15):1211. doi:10.3390/cells14151211

28. Habibey R, Striebel J, Schmieder F, Czarske J, Busskamp V. Long-term morphological and functional dynamics of human stem cell-derived neuronal networks on high-density micro-electrode arrays. Front Neurosci. 2022;16:951964. doi:10.3389/fnins.2022.951964

29. Parodi G, Zanini G, Collo L, et al. In vitro electrophysiological drug testing on neuronal networks derived from human induced pluripotent stem cells. Stem Cell Res Ther. 2024;15(1):433. doi:10.1186/s13287-024-04018-2

30. Sit TPH, Feord RC, Dunn AWE, et al. MEA-NAP: A flexible network analysis pipeline for neuronal 2D and 3D organoid multielectrode recordings. Cell Rep Methods. 2024;4(11):100901. doi:10.1016/j.crmeth.2024.100901

31. Escoubas CC, Guney E, Miravet ÀT, et al. NOVA: a novel R-package enabling multi-parameter analysis and visualization of neural activity in MEA recordings. bioRxiv: The Preprint Server for Biology. Preprint posted online October 2, 2025:2025.10.01.679841. doi:10.1101/2025.10.01.679841

32. Biffi E, Menegon A, Regalia G, Maida S, Ferrigno G, Pedrocchi A. A new cross-correlation algorithm for the analysis of “in vitro” neuronal network activity aimed at pharmacological studies. J Neurosci Methods. 2011;199(2):321–327. doi:10.1016/j.jneumeth.2011.05.010

33. Teppola H, Aćimović J, Linne ML. Unique Features of Network Bursts Emerge From the Complex Interplay of Excitatory and Inhibitory Receptors in Rat Neocortical Networks. Front Cell Neurosci. 2019;13. doi:10.3389/fncel.2019.00377

34. Park S, Cho SJ, Kim CY, Chung HM, Lee SG. Non-Invasive electrophysiological monitoring of cardiac organoids using 3D-Net-assisted microelectrodes array platform. Sci Rep. 2026;16(1):4428. doi:10.1038/s41598-025-34504-3

35. Weiser-Bitoun I, Mazgaoker S, Assayag S, Davoodi M, Alexandrovich A, Yaniv Y. PhysioMEA: Signal processing platform for rate and rhythm analysis of multi-electrode array cardiac electrophysiological recordings. J Mol Cell Cardiol. 2026;210:137–149. doi:10.1016/j.yjmcc.2025.11.006

36. Hausdorff WP, Caron MG, Lefkowitz RJ. Turning off the signal: desensitization of β-adrenergic receptor function. FASEB J. 1990;4(11):2881–2889. doi:10.1096/fasebj.4.11.2165947

37. Yu SS, Lefkowitz RJ, Hausdorff WP. Beta-adrenergic receptor sequestration. A potential mechanism of receptor resensitization. J Biol Chem. 1993;268(1):337–341. doi:10.1016/S0021-9258(18)54155-7

38. Limas CJ, Limas C. Rapid recovery of cardiac beta-adrenergic receptors after isoproterenol-induced “down”-regulation. Circ Res. 1984;55(4):524–531. doi:10.1161/01.RES.55.4.524

